# Harbingers of Aggressive Prostate Cancer: Precision Oncology Case Studies

**DOI:** 10.1101/2021.11.30.468210

**Authors:** Joanna Cyrta, Davide Prandi, Arshi Arora, Daniel H. Hovelson, Andrea Sboner, Antonio Rodriguez, Tarcisio Fedrizzi, Himisha Beltran, Dan R. Robinson, Anurandha Gopalan, Lawrence True, Peter S. Nelson, Brian D. Robinson, Juan Miguel Mosquera, Scott A. Tomlins, Ronglai Shen, Francesca Demichelis, Mark A. Rubin

## Abstract

Primary prostate cancer (PCa) can show marked molecular heterogeneity. However, systematic analyses comparing primary PCa and matched metastases in individual patients are lacking. We aimed to address the molecular aspects of metastatic progression while accounting for heterogeneity of primary PCa.

In this pilot study, we collected 12 radical prostatectomy (RP) specimens from men who subsequently developed metastatic castration-resistant prostate cancer (mCRPC). We used histomorphology (Gleason grade, focus size, stage) and immunohistochemistry (IHC) (ERG and p53) to identify independent tumors and/or distinct subclones of primary PCa. We then compared molecular profiles of these primary PCa areas to matched metastatic samples using whole exome sequencing (WES) and amplicon-based DNA and RNA sequencing. Based on combined pathology and molecular analysis, seven (58%) RP specimens harbored monoclonal and topographically continuous disease, albeit with some degree of intra-tumor heterogeneity; four (33%) specimens showed true multifocal disease; and one displayed monoclonal disease with discontinuous topography. Early (truncal) events in primary PCa included *SPOP* p.F133V (one patient), *BRAF* p.K601E (one patient), and *TMPRSS2*:ETS rearrangements (nine patients). Activating *AR* alterations were seen in eight (67%) mCRPC patients, but not in matched primary PCa. Hotspot *TP53* mutations, found in metastases from three patients, were readily present in matched primary disease. Alterations in genes encoding epigenetic modifiers were observed in several patients (either shared between primary foci and metastases or in metastatic samples only).

WES-based phylogenetic reconstruction and/or clonality scores were consistent with the index focus designated by pathology review in six out of nine (67%) cases. The three instances of discordance pertained to monoclonal, topographically continuous tumors, which would have been considered as unique disease in routine practice.

Overall, our results emphasize pathologic and molecular heterogeneity of primary PCa, and suggest that comprehensive IHC-assisted pathology review and genomic analysis are highly concordant in nominating the “index” primary PCa area.

## Introduction

Primary prostate cancer (PCa) is a multifocal disease in up to 80% of PCa patients [1, 2]. In addition, it has been shown to display marked inter- and intra-tumor heterogeneity at the genomic, transcriptomic and DNA methylation levels [3-6]. Conversely, metastatic castration-resistant prostate cancer (mCRPC) appears to be of clonal origin, even though it can acquire molecular heterogeneity through subclonal evolution [4, 7]. There is a critical knowledge gap regarding the molecular mediators of PCa progression to mCRPC in individual patients. While pathology criteria associated with aggressive disease, e.g. Gleason grade or focus size, are typically used to nominate the dominant (index) focus in radical prostatectomy (RP) specimens, the validity of this approach has not been addressed by systematic studies, and has even been challenged by occasional case reports [8].

Due to the long time between primary therapy and development of metastatic disease, only a few studies have analyzed matched primary and metastatic samples. As part of two precision oncology trials, the CRPC500 [9, 10] and the Weill Cornell Medicine (WCM) Precision Oncology cohort [11], we present a comprehensive pathology and genomic analysis of RP specimens from 12 men who subsequently developed mCRPC. We compare the genomics of multiple areas of primary PCa with matched metastatic disease, with the aim of nominating the index focus and identifying the histopathological and molecular criteria that could be associated with metastatic outcome.

## Materials and methods

### Sample collection and pathology review

Medical records of patients with metastatic, castration-resistant prostate cancer (mCRPC) enrolled in WCM precision cancer care program and/or in the Stand-Up-To-Cancer (SU2C) CRPC500 study [10] were interrogated to identify patients who had previously undergone radical prostatectomy (RP). Only patients for whom complete RP pathology material (all H&E slides and blocks) could be retrieved were included. Three pathologists (J.M.M., B.R. and J.C.) jointly reviewed all H&E slides for each RP specimen. Immunohistochemistry (IHC) for ERG and p53 was performed on all tumor areas. Potentially distinct areas (i.e., either subclones of the same primary disease or truly independent tumors) were selected for sequencing based on topography (i.e., discontinuous nature of tumor areas), individual Gleason score, histomorphology, and/or IHC. The putative index focus was nominated based on the highest individual Gleason score (primary criterion) and focus size (secondary criterion). Seminal vesicle invasion and metastatic regional lymph nodes removed at time of RP were also included in this evaluation and by principle, considered as putative index disease. Pathology review of metastatic samples (frozen sections) was previously performed as part of the WCM precision cancer care and/or the SU2C CRPC500 study [10]. Both studies were performed with appropriate written patient consent and IRB approval.

### Immunohistochemistry

Immunohistochemistry (IHC) was done on sections of FFPE tissue using a Bond III automated immunostainer and the Bond Polymer Refine detection system (Leica Microsystems, IL, USA), with the following antibodies and conditions (dilution, heat-mediated antigen retrieval solution, retrieval time): anti-ERG (Abcam, clone EPR386, 1/100, H1, 30min); anti-p53 (Cell Signaling Technologies, clone DO-7, 1/100, H1, 20min).

### Whole-exome sequencing

WES was performed using previously validated WCM protocols [12]. Briefly, DNA was extracted from macrodissected unstained slides of FFPE tissue cut at 10um (for each primary tumor focus); from cored frozen, OCT-embedded tissue (for metastases); and from peripheral blood lymphocytes (for germline control). DNA extraction was done using Promega Maxwell 16 MDx (Promega, Madison, WI). DNA quality was confirmed by real-time PCR using Bioanalyzer (Agilent Technologies, Santa Clara, CA). Libraries were prepared using exome capture of 21,522 genes with the HaloPlex System (Agilent). Sequencing was performed on Illumina HiSeq 2500 (San Diego, CA) in 100bp paired-end mode [12]. Reads were aligned to GRC37/hg19 reference using BWA [13] and processed accordingly to Whole Exome Sequencing Test for Cancer – ExaCT-1 - pipeline v0.9 [12].

To identify SNVs, we applied both MuTect [14] and SNVseeqer [15] to nominate putative aberrant genomic positions. Identified genomic positions were filtered requiring coverage of at least 10 reads, a read count of the alternative base of at least 3, and a minimum variant allelic fraction (VAF) of 5%. Filtered positions were inspected with ASEQ [16], which provides a read count for each of the four bases in tumor and matched normal samples. Genomic positions where the read count of the alternative base in normal matched sample was greater than 0 were considered not aberrant. The list of aberrant genes was divided into 3 tiers: tier1 containing genes in the EXaCT-2 WCM test, tier2 containing genes in COSMIC Cancer Gene Census, tier3 containing all remaining genes.

For metastatic samples for Patients 11 and 12, WES data from fresh-frozen tissue were obtained through the CRPC500 study and generated as previously described [9].

### Targeted DNA and RNA sequencing

DNA and RNA extraction, amplicon-based next generation sequencing, and bioinformatics analysis were performed as previously described [5].

### Phylogenetic reconstruction and clonality analysis

Based on SNV calls, primary and metastatic samples were analyzed for clonal relatedness through phylogenetic tree reconstruction and clonality score. The final list of aberrant genomic positions was composed of: i) mutations in genes in tier1 or tier2 called by either Mutect or SNVseeqer, and ii) mutations in genes in tier3 called by both Mutect and SNVseeqer. Maximum-parsimony trees were inferred using the phangorn package [17]. The germline sample was designated as the root. Primary and metastatic tumor samples were plotted as descendent nodes, with edge lengths indicating the number of alterations newly accumulated in descendant nodes. Samples sharing common mutations formed a node. Clonality scores were determined as previously described [18] by performing conditional test on sites with mutation in one or both tumors. Marginal probabilities at each locus were estimated via empirical relative frequencies from the somatic mutations data set from publicly available TCGA PRAD study. [19].

## Results

### Pathology and clinical findings

Twelve patients were included in this pilot cohort (nine enrolled in WCM precision cancer care and three in the SU2C CRPC500 study [10]). The number and type of samples per patient, pathology results and IHC are summarized in **Figure 1** and in **Supplementary table ST1**. We assigned individual Gleason scores to each discrete sample. Samples showed a range from Gleason score 6 (3+3) to 9 (5+4). Disease stage per patient ranged from pT2aN0 to pT3bN1. Partial neuroendocrine features (positive synaptophysin expression with maintained AR expression and an adenocarcinoma morphology) were noted in primary PCa from one patient (Patient 7) and in the metastasis of another patient (Patient 6). Neuroendocrine prostatic carcinoma (NEPC) with adequate morphology [20] and loss of AR expression was seen in the RP specimen from one patient (Patient 12) and in metastases from two patients (Patients 12 and 7). ERG expression was seen in all or a subset of samples in 8 (67%) patients. The time from prostatectomy to metastatic sample collection (per sample) ranged from 15.9 to 230.5 months (mean, 47.1 months). Metastatic sites included bone, lymph node, liver and brain. Initial pathology review of RP specimens, including IHC results for ERG and p53, but blinded to the genomic data, interpreted eight (67%) RP specimens as monoclonal disease with intra-tumor heterogeneity and four (33%) RP specimens as harboring true multiclonal disease (Patients 3, 6, 11 and 12). In three of the multiclonal cases, the additional tumors were only small-volume Gleason grade 6 foci (Patients 3, 6 and 11). In one case (Patient 12), as many as six potentially independent tumors were identified, including three Gleason grade 7 and three Gleason grade 6 lesions (**Figure 2**). Using Gleason grade, tumor size and stage, one or multiple index tumor areas were nominated by pathology for all cases.

**Figure 1.**
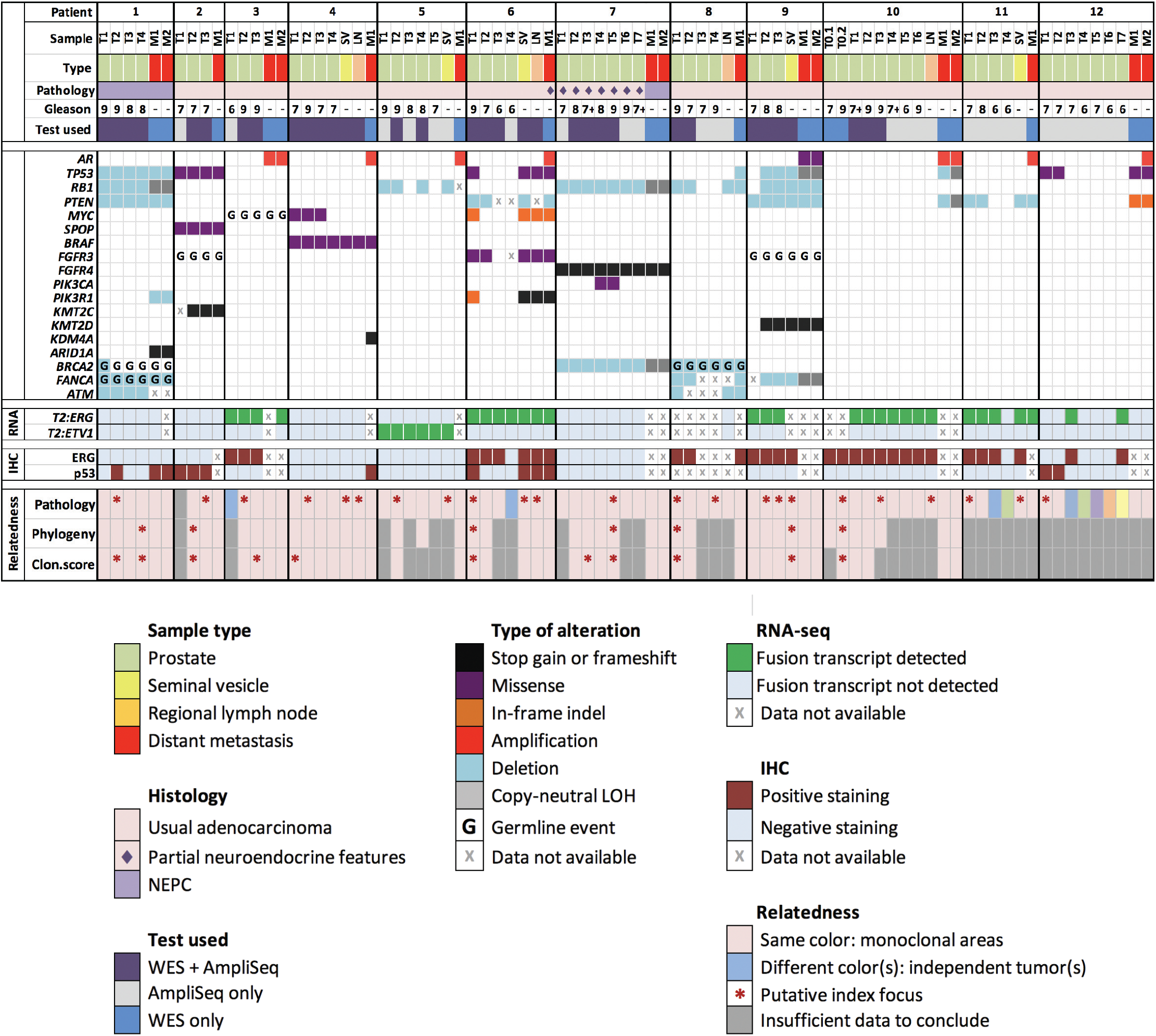
A summary of all analyzed samples, including their histopathology characteristics, a schematic representation of selected genomic findings, immunohistochemistry (IHC) results, and clonal relatedness as interpreted by pathology and by genomics, i.e. phylogenetic reconstruction (“Phylogeny”) and clonality score (“Clon.score”); 7+ indicates Gleason score 7 with tertiary 5 pattern; *T2:ERG* stands for *TMPRSS2:ERG*; *T2:ETV1* stands for *TMPRSS2:ETV1*.

**Figure 2.**
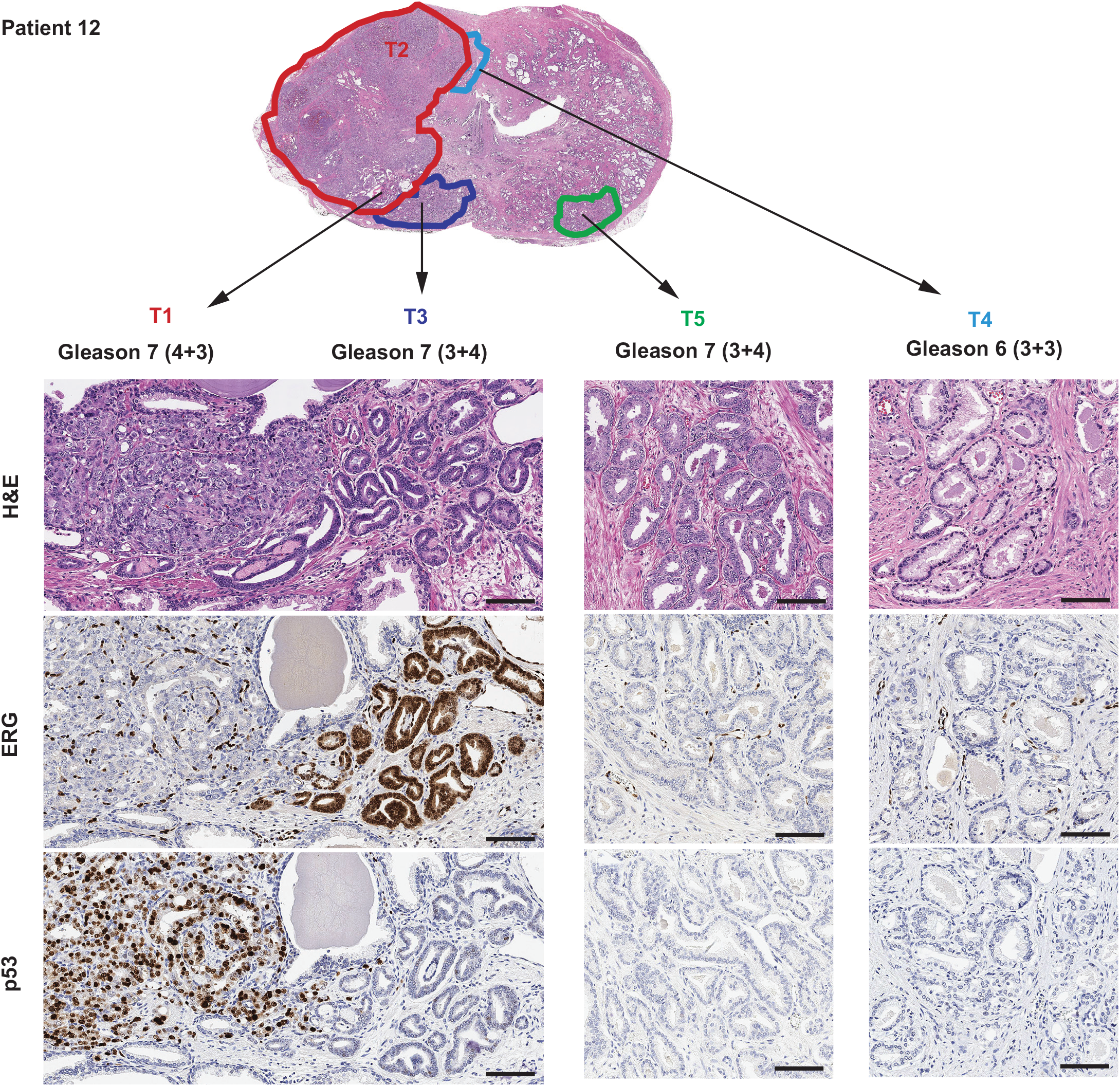
An example of pathology annotation in a case of multiclonal prostate cancer (Patient 12), including collision tumors. Pathology review and immunohistochemistry (IHC) identified 6 potentially independent tumors: a large, Gleason grade 7 focus which was ERG-/p53+ by IHC (samples T1 and T2); a Gleason grade 7 focus, which was ERG+/p53-by IHC (sample T3) and in collision with T1; a topographically distinct Gleason grade 7 focus which was ERG-/p53-by IHC (sample T5); and 3 additional low-volume distinct Gleason grade 6 foci (sample T4, pictured, and samples T6 and T7, not shown). Topographical confluence (tumor collision) was noted between areas T1 and T3, and between areas T2 and T4.

### Genomic landscape of primary PCa and paired metastatic samples

WES was performed on 36 primary tumor areas from 10 patients (2-6 samples per patient) (**Supplementary tables ST2-ST4, Figures 3 and 4, Supplementary Fig.S1-S8)**. Targeted DNA sequencing was performed on 63 primary samples from 12 patients (3-7 per patient) (**Supplementary tables ST5-ST6)**. Results were compared to metastatic tumor samples (18 samples from 12 patients, 1-2 samples per patient), which previously underwent WES [10, 11]. The most frequent genomic event found in the metastases, and not observed in the matching primary samples, were *AR* activating alterations, seen in 8 (67%) patients (amplification in seven cases, activating point mutations in one case). The second most frequent alteration type found in metastases were *TP53* missense hotspot mutations, identified in three (25%) patients. Importantly, in contrast to *AR* activating alterations, matched analysis confirmed that these *TP53* alterations were present in some (two cases) or all (one case) areas of primary PCa in each patient (**Figures 1-3**).

**Figure 3.**
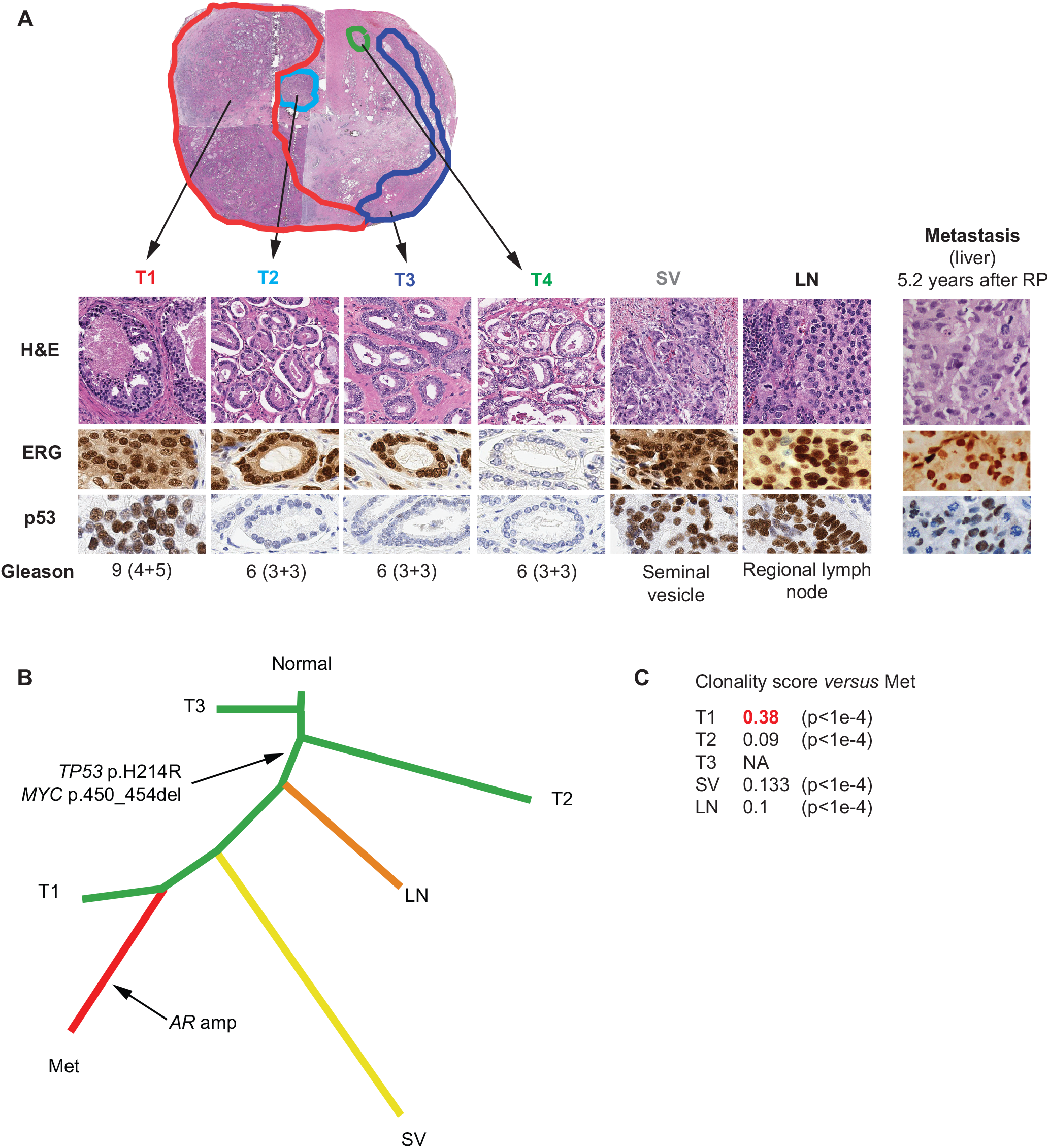
A summary of pathology features, phylogeny and clonality analyses for Patient 6. **A**. Pathology features of each sample, including histomorphology, ERG and p53 immunohistochemistry (IHC). Tumor focus 1 (T1) is the dominant (index) focus with highest Gleason grade and size. The T4 focus has a different ERG IHC status from other foci, and was thus regarded as an independent tumor on pathology evaluation. Aberrant p53 immunostaining is shared by T1, seminal vesicle (SV), lymph node (LN) and metastasis (Met). H&E: hematoxylin-eosin. **B**. Phylogenetic reconstruction using 79 genes and 110 events confirms that samples T1, SV and LN are more closely related to the metastasis than samples T2 or T3, as evidenced by a common “trunk” which includes the *TP53* mutation. **C**. A summary of clonality score results. Sample T1 shows the highest clonality score (highlighted in red) with respect to the metastatic sample.

**Figure 4.**
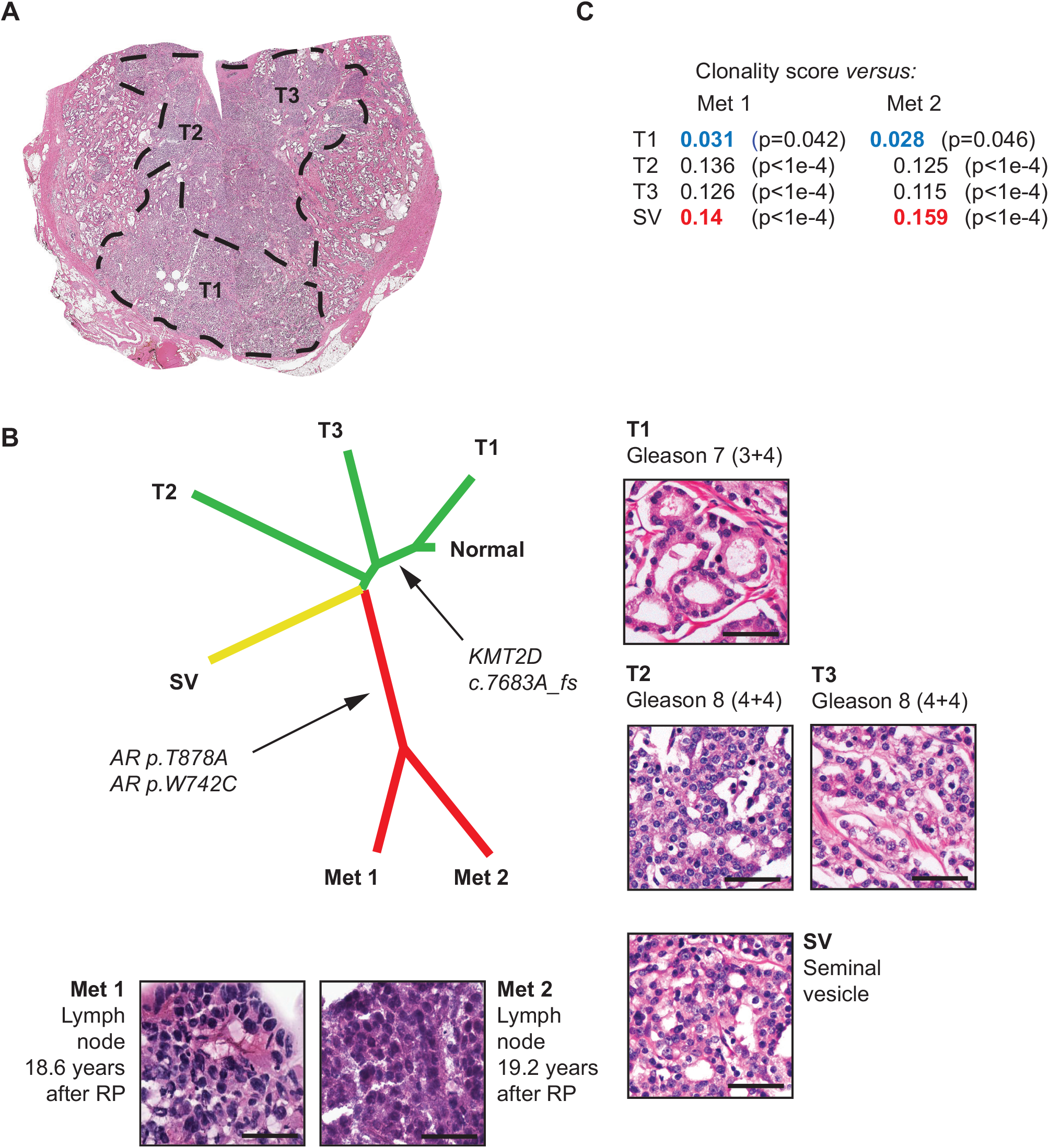
A summary of pathology features, phylogeny and clonality analyses for Patient 9. **A**. Low power pathology view. This case was interpreted as a unifocal, monoclonal tumor, albeit with the T1 tumor area showing a lower Gleason grade than areas T2 and T3. **B**. Phylogenetic reconstruction using 71 genes and 103 events, confirming that samples T2, T3 and seminal vesicle (SV) are more closely related to the metastasis than T1. Detailed histomorphology for all foci is also shown. **C**. A summary of clonality score results. Sample SV shows the highest clonality score (highlighted in red) with respect to both metastatic samples. Note that sample T1 (with the lowest individual Gleason score) also shows the lowest clonality score (highlighted in blue).

Comparative analysis of paired samples from individual patients identified some early (“truncal”) events, present at high allelic frequencies in all primary samples and in the metastases. These included: a hotspot p.F133V *SPOP* mutation in Patient 2, and a *BRAF* p.K601E mutation in Patient 4. In addition, RNA sequencing identified *TMPRSS2:ERG* and *TMPRSS2:ETV1* fusion transcripts in 6 patients and in one patient, respectively. These fusion transcripts were detected across multiple samples and, when data were available, in the matched metastasis. The *TMPRSS2:ERG* status was consistent with ERG IHC in all but one sample. This one sample (Patient 6, T4) was ERG-negative on IHC and displayed a relatively low *TMPRSS2:ERG* signal on RNA-seq, likely due to contamination with cells from the nearby ERG-positive tumor focus T3 (**Supplementary Fig.S9**).

Several cases harbored alterations in genes encoding epigenetic modifiers. *KMT2D (MLL2/4)* c.7683A_fs in Patient 9 was shared by all primary samples, except for one area with a lower Gleason score, and by the two metastases (**Figure 4**). Other alterations included: *KMT2C (MLL3)* p.C1013* in Patient 2, shared by all primary samples and the metastasis (**Supplementary Fig.S2**); *KDM4A* (*JMJD2*) p.L696_fs in Patient 4, detected in the metastasis only (**Supplementary Fig.S4**); and p.1479_fs in *ARID1A*, encoding a member of the SWI/SNF complex, detected in both metastatic samples in Patient 1, but not in the primary samples (**Supplementary Fig.S1**).

*BRCA2* germline mutations were found in two (16.7%) cases (Patients 1 and 8), one of which showed neuroendocrine features in the primary and metastatic tumors.

### Assessing clonal relatedness between primary and metastatic samples

Phylogenetic reconstruction using WES SNV calls allowed to nominate an index focus/foci in seven (70%) of the 10 patients for whom WES on the primary samples was available, and clonality scores could be obtained for nine patients (90%) (**Figure 1, Supplementary table ST1**). Both approaches showed highly concordant results.

In six of the nine cases (67%), the index focus nominated by phylogenetic reconstruction and/or by the highest clonality score was consistent with the one proposed by pathology review.

Discordance was seen in three cases (Patients 2, 3 and 4), all of which were monoclonal tumors with continuous spread.

In addition, although phylogeny and clonality analyses were not available for Patient 12, concordance between pathology and genomics could be inferred owing to the presence of a shared *TP53* mutation (p.R248W) and the absence of a *TMPRSS2:ERG* fusion in the index focus and the metastases (**Figure 1**).

### Gene expression analysis

Targeted RNA sequencing was performed on 58 primary and 5 metastatic samples (**Figure 5**). Unsupervised hierarchical clustering using gene expression levels from RNA-seq further supported molecular similarity between genomically similar PCa areas, especially in monoclonal primary tumors. Conversely, sample clustering results for Patients 3, 6, 11 and 12 were potentially consistent with multiclonality. Of note, mCRPC samples from four patients (two bone and two liver metastases) formed a distinct transcriptomic cluster, characterized by high expression of proliferation genes (**Figure 5**, box), consistent with results using this approach on a separate comependium of localized treatment-naive PCa and CRPC metastases [5].

**Figure 5.**
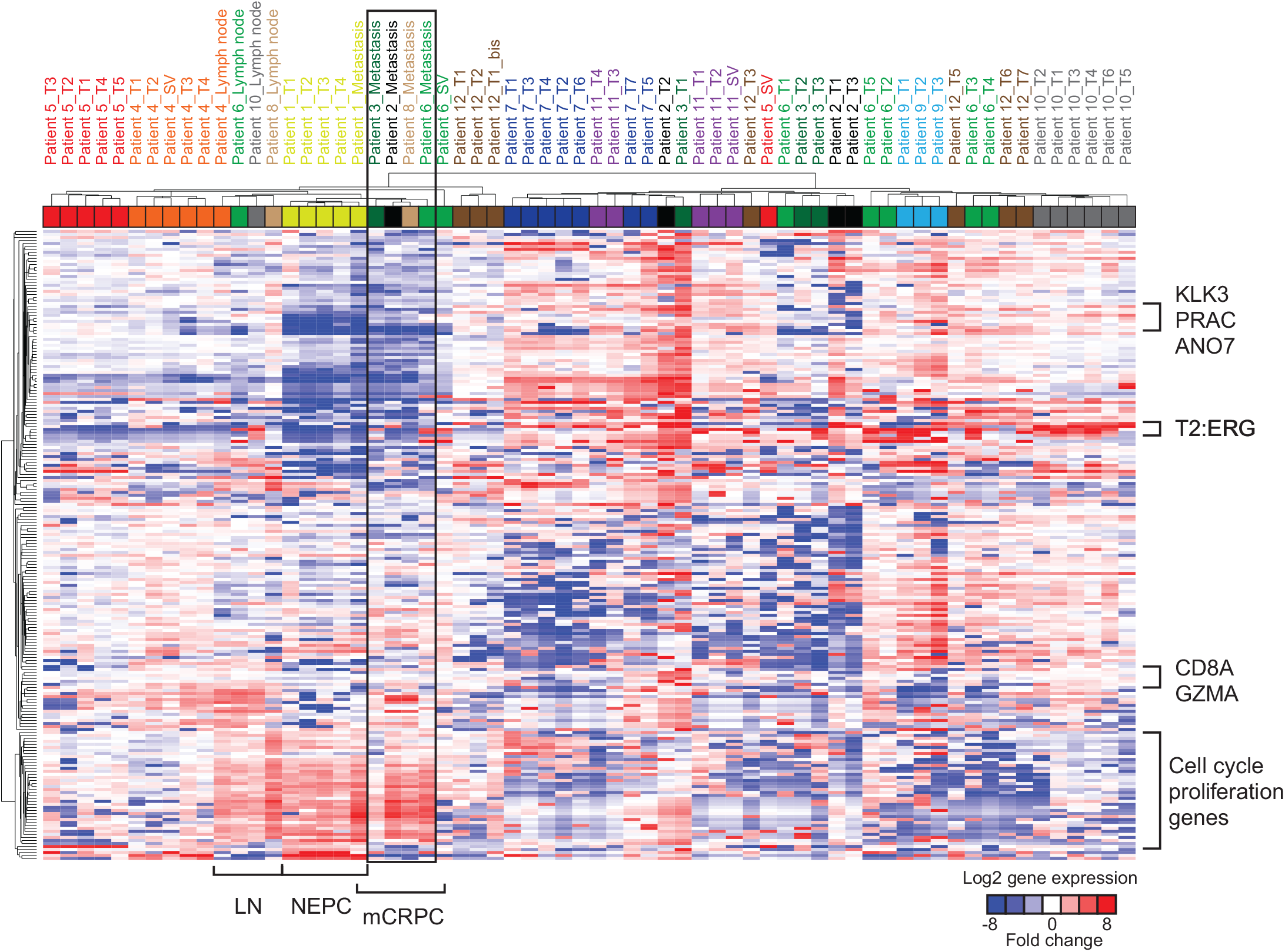
Unsupervised sample clustering and heatmap representation of RNA-seq gene expression levels using 204 target expression amplicons. Only samples that have passed quality control as detailed in Salami et al. [5] are shown. T2:ERG: *TMPRSS2:ERG* fusion transcripts, LN: regional lymph node, NEPC: neuroendocrine prostatic carcinoma, mCRPC: metastatic castration-resistant prostatic carcinoma. The black box indicates mCRPC samples from four patients.

## Discussion

### Genomic heterogeneity of PCa and different scenarios for PCa clonality

Primary PCa has previously been shown to represent a molecularly heterogeneous disease [3-6]. When considering clonality of primary PCa, four scenarios can be proposed, all of which were encountered in our pilot cohort (**Figure 6, Supplementary table ST1**).

**Figure 6.**
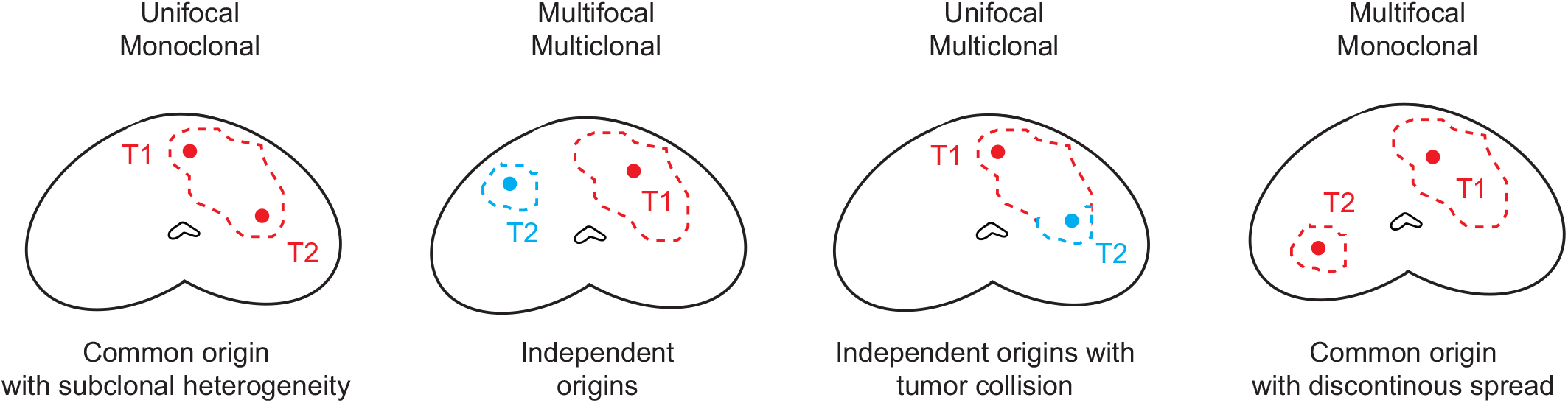
Possible scenarios for clonality interpretation of primary prostate cancer.

In one scenario, tumors are monoclonal (all areas have a common origin) and show topographically continuous spread, but still demonstrate intra-tumor heterogeneity due to emergence of distinct subclones [3, 4]. This was the most common scenario in our study, seen in seven (58%) cases.

Another possibility is the presence of true multiclonal tumors with independent origins in the RP specimen [21-24]. Only four (33.3%) cases from our cohort showed true multiclonal disease. The rate of true multiclonal disease in our study was lower than the 50-80% previously reported in large RP series [1, 2]. This could be due to selection bias, as we only included patients who subsequently developed metastases, thus selecting for aggressive primary disease, which may potentially override synchronous low-volume lesions or not leave the time for such lesions to develop. This is in keeping with a study by Wise et al., who found that a greater number of “lesser cancers” correlates with a smaller volume of the index cancer [2]. In most instances, and in three (25%) of our patients, such additional foci are low-grade and low-volume (<0.5 cm^3^), and represent an incidental finding rather than clinically relevant disease [1, 2, 25]. Critically, these multiclonal tumors may either be topographically distinct (multifocal), which represents the second scenario, or confluent (collision tumors), which represents the third scenario (**Figures 2 and 5**). We observed collision of clonally independent tumors, confirmed by discordant ERG IHC status, in three RP specimens (Patients 6, 11, and 12). This phenomenon of collision tumors suggests that topography alone is not reliable in determining relatedness of PCa areas. Lastly, the disease may have a common clonal origin, but discontinuous topography (monoclonal-multifocal), potentially due to a particular form of tumor dissemination or to RP specimen processing errors; however, contrarily to what has been proposed by some authors [26], this appears to be a rare scenario (one patient in our study).

Overall, some degree of molecular heterogeneity of PCa (either in a “subclonal” or “multiclonal” form) was observed in all cases in our cohort. This suggests that random sampling of a single primary tumor area may not be sufficient to capture the entire genomic landscape of a patient’s primary disease [5].

In our study, the index tumor focus nominated by pathology in RP specimens was concordant with the one nominated by genomics in most cases where a conclusion could be reached. The three instances of discordance (Patients 2, 3, and 4) pertained to monoclonal and topographically continuous tumor areas (**Supplementary Fig.S2-4**), which would have been considered as a unique tumor in routine pathology practice.

In contrast to our findings, Haffner et al. reported a case in which a tumor area with Gleason pattern 3 and organ confined, and not the areas with predominant pattern 4, was found to be most closely related to the metastasis [8]. However, this was not an independent low-volume Gleason grade 6 tumor, but an area within a larger lesion showing various proportions of patterns 3 and 4. In addition, this specific area also harbored a *TP53* mutation.

### Comparative analysis of primary PCa and matched metastases

Paired analysis of multiple primary samples and matched metastases from individual patients is particularly helpful for determining at which point of disease progression genomic alterations of interest appear. Activating *AR* alterations were found only in metastatic samples, in agreement with the previously reported major enrichment of such alterations in metastatic cohorts [7, 9, 27-31], although exceptional instances of *AR* alterations in primary disease have been reported [7]. This is likely due to the fact that metastatic samples from our study represented castration-resistant disease with AR alterations driving treatment resistance [9, 30]. The metastatic NEPC in Patients 1 and 7 did not show *AR* amplification, consistent with “indifference” of NEPC to AR signaling [29].

In addition, we were able to demonstrate in individual patients that some alterations reported to be enriched in metastatic cohorts readily arise in the primary disease. In particular, *TP53* alterations, which are more frequent in metastatic PCa cohorts compared to primary PCa cohorts [9, 30-32], were found in metastatic samples from three patients in our study. In all three patients, these alterations were also detected in the primary cancer, and were subclonal in two cases. This could warrant further assessment of *TP53* alterations as a predictive factor for metastatic relapse in patients with localized PCa, and appears consistent with the case reported by Haffner et al. [8].

Conversely to subclonal *TP53* mutations, some alterations seem to represent early (“truncal”) oncogenic driver events, as they were clonally present throughout all samples from the primary disease and also shared by the metastases: *SPOP* p.F133V (a well described hotspot mutation in PCa [33, 34]), *BRAF* p.K601E (a potentially activating variant, which has previously been reported in PCa [19]), and *TMPRSS2:ERG* or *ETV1* rearrangements, in keeping with the early and clonal nature of this alteration [35]. Although to date, there are no therapies targeting these specific alterations, these examples suggest that in cases where the genomic assessment of the metastatic disease is not possible, sequencing of the primary tumor may still provide valuable information on driver mutations.

Our results also highlight the potential role of epigenetic processes in metastatic PCa progression. Multiple studies have found alterations in *KMT2C* and *KMT2D*, which encode histone methyltransferases, to be enriched in metastatic PCa cohorts as compared to primary PCa [7, 9, 30, 31, 36]. Similar to *TP53*, through paired analysis, we showed that truncating *KMT2C* and *KMT2D* alterations detected in the metastases (Patients 2 and 9, respectively) were already present in the matched primaries. It could be important to investigate in larger cohorts whether the presence of such alterations in primary PCa is a risk factor for metastatic recurrence. Alterations in *KDM4A* and *ARID1A* could also warrant further investigation [19, 30, 36].

The frequency of pathogenic *BRCA2* germline alterations in mCRPC patients has previously been reported at 5.3% [37]. In our cohort, germline *BRCA2* alterations were found in two patients (16.7%). In one patient (Patient 1), the primary tumor and the metastases were NEPC. Because of the limited size of this pilot cohort, findings from these “n-of-1” cases need to be confirmed in larger studies. Nevertheless, collecting archival RP specimens from patients with mCRPC is exceedingly difficult, as the period of time between primary therapy and metastatic disease can be very long (up to 18 years in our cohort, Patient 8). Another important hurdle in analyzing genomic relatedness between multiple samples are variations in tumor DNA content. Because of this caveat, copy number alterations could not be included in the phylogeny analyses, and SNV-based analyses may have been negatively impacted for some samples (e.g. regional lymph node samples) in which tumor purity was typically low. Lastly, other IHC biomarkers previously used in assessing PCa heterogeneity, such as PTEN and SPINK1 [38], could potentially be employed in future studies to even better discriminate tumor populations on pathology review.

## Conclusion

Our study confirms that primary PCa is characterized by marked pathologic and genomic heterogeneity, which must be considered when selecting tumor areas for precision medicine studies. This is one of the first studies to formally address the still controversial concept of the “index lesion” in primary PCa. Our findings in this pilot cohort show high concordance between a comprehensive, IHC-assisted pathology review and genomic analyses for nominating the “index focus”, although the specificity of these approaches should be validated in larger cohorts.

## Acknowledgements

We thank patients and their families for participating in genomics, transcriptomics, and precision cancer care studies.

We are thankful for expert assistance from the Translational Research Program at WCM Pathology and Laboratory Medicine (Bing He, Leticia Dizon, Yifang Liu, Mai Ho) and the WCM CLC Genomics Core Facility (Jenny Xiang). We are grateful to Makayla DeJong (Seattle Cancer Care Alliance) for her help with retrieving archival pathology material. We thank Dr. Christopher Barbieri (Weil Cornell Medicine) for his valuable suggestions regarding this study. We acknowledge expert assistance from Mariana Ricca at the University of Bern in preparing the manuscript for submission.

## Data Sharing Policy

Data are available for bona fide researchers who request it from the authors.

## Supplementary Figure legends

**Supplementary figure S1**. A summary of pathology features, phylogene and clonality analyses for Patient 1. **A**. Phylogenetic reconstruction using 135 genes and 274 events, and pathology features corresponding to each focus. Histomorphology and p53 immunostaining are shown. Note that despite diffuse nuclear staining for p53 in some samples, no *TP53* mutation was detected in these samples with the methods used herein. **B**. A schematic heatmap representation of the most significant SNV events used in the phylogenetic reconstruction across samples. **C**. A summary of clonality score results. Note that the clonality score is highest for the T1 focus with respect to the Metastasis 1 sample, and in the T2 focus (predicted as index focus by pathology) with respect to the Metastasis 2 sample. RP: radical prostatectomy; H&E: hematoxylin-eosin. Scale bars: 50 μm.

**Supplementary figure S2**. A summary of pathology features, phylogeny and clonality analyses for Patient 2. **A**. Phylogenetic reconstruction using 37 genes and 47 events and pathology features corresponding to each focus. Histomorphology and p53 immunostaining are shown. **B**. A schematic heatmap representation of SNV events used in the phylogenetic reconstruction across samples. **C**. A summary of clonality score results. RP: radical prostatectomy; ADT: androgen deprivation therapy; H&E: hematoxylin-eosin. Scale bars: 50 μm.

**Supplementary figure S3**. A summary of pathology features, phylogeny and clonality analyses for Patient 3. **A**. Phylogenetic reconstruction using 50 genes and 64 events and pathology features corresponding to each focus. Histomorphology and ERG immunostaining are shown. **B**. A schematic heatmap representation of the most significant SNV events used in the phylogenetic reconstruction across samples. **C**. A summary of clonality score results. RP: radical prostatectomy; ADT: androgen deprivation therapy; H&E: hematoxylin-eosin.

**Supplementary figure S4**. A summary of pathology features, phylogeny and clonality analyses for Patient 4. **A**. Phylogenetic reconstruction using 87 genes and 142 events and pathology features corresponding to each focus. **B**. A schematic heatmap representation of the most significant SNV events used in the phylogenetic reconstruction across samples. **C**. A summary of clonality score results. RP: radical prostatectomy; ADT: androgen deprivation therapy. Scale bars: 50 μm.

**Supplementary figure S5**. A summary of pathology features, phylogeny and clonality analyses for Patient 5. **A**. Phylogenetic reconstruction using 10 genes and 15 events and pathology features corresponding to each focus. **B**. A schematic heatmap representation of SNV events used in the phylogenetic reconstruction across samples. The clonality scores could not be computed for this case. RP: radical prostatectomy; N/A: not available.

**Supplementary figure S6**. A summary of pathology features, phylogeny and clonality analyses for Patient 7. **A**. Phylogenetic reconstruction using 111 genes and 145 events and pathology features corresponding to each focus. **B**. A schematic heatmap representation of the most significant SNV events used in the phylogenetic reconstruction across samples. **C**. A summary of clonality score results. RP: radical prostatectomy.

**Supplementary figure S7**. A summary of pathology features, phylogeny and clonality analyses for Patient 8. **A**. Phylogenetic reconstruction using 10 genes and 15 events and a schematic representation of the two tumor foci. **B**. A schematic heatmap representation of SNV events used in the phylogenetic reconstruction across samples. **C**. A summary of clonality score results.

**Supplementary figure S8**. A summary of pathology features, phylogeny and clonality analyses for Patient 10. **A**. Phylogenetic reconstruction using 22 genes and 30 events and pathology features corresponding to each focus. **B**. A schematic heatmap representation of SNV events used in the phylogenetic reconstruction across samples. **C**. A summary of clonality score results.

**Supplementary figure S9**. Patient 6, primary tumor focus T4. This focus showed some *TMPRSS2-ERG* signal on RNA-seq, but was negative for ERG by IHC. The topographic proximity of the T3 focus (ERG-positive) and contamination during the macrodissection step may have accounted for this discrepancy. H&E: hematoxylin eosin, IHC: immunohistochemistry.

## Supplementary Tables legends

**ST1**: Sample summary: A summary of samples for each patient, together with pathology findings and interpretation of clonality based on pathology and on genomics.

**ST2**: WES SNV-clonality: A list of WES-based SNV calls per sample, as used for phylogeny and clonality analyses (after applying additional filters described in Methods)

**ST3**: WES indels: WES-based short insertions and deletions per sample.

**ST4**: WES CNA: WES-based copy number alterations (CNA) per sample. Only selected genes and only samples for which allele-specific CNA could be computed are shown.

**ST5**: NGS SNV: Targeted sequencing-based SNV and indel calls per sample.

**ST6**: NGS CNA: Targeted sequencing-based CNA calls per sample.

