## Supplemental Figures for "Harbingers of Aggressive Prostate Cancer: Precision Oncology Case Studies"

### Supplementary figure S1

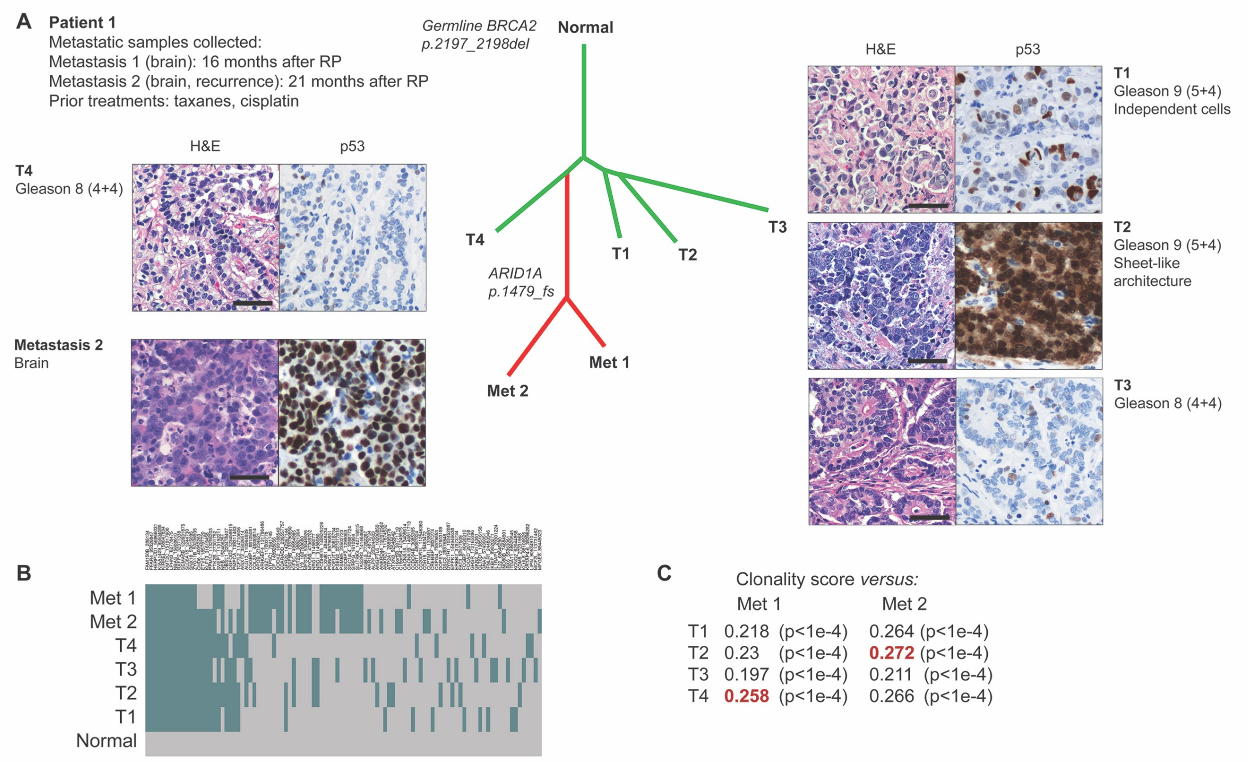


**Supplementary figure S1.** A summary of pathology features, phylogene and clonality analyses for Patient 1. **A.** Phylogenetic reconstruction using 135 genes and 274 events, and pathology features corresponding to each focus. Histomorphology and p53 immunostaining are shown. Note that despite diffuse nuclear staining for p53 in some samples, no *TP53* mutation was detected in these samples with the methods used herein. **B.** A schematic heatmap representation of the most significant SNV events used in the phylogenetic reconstruction across samples. **C.** A summary of clonality score results. Note that the clonality score is highest for the T1 focus with respect to the Metastasis 1 sample, and in the T2 focus (predicted as index focus by pathology) with respect to the Metastasis 2 sample. RP: radical prostatectomy; H&E: hematoxylin-eosin. Scale bars: 50 μm.

### Supplementary figure S2

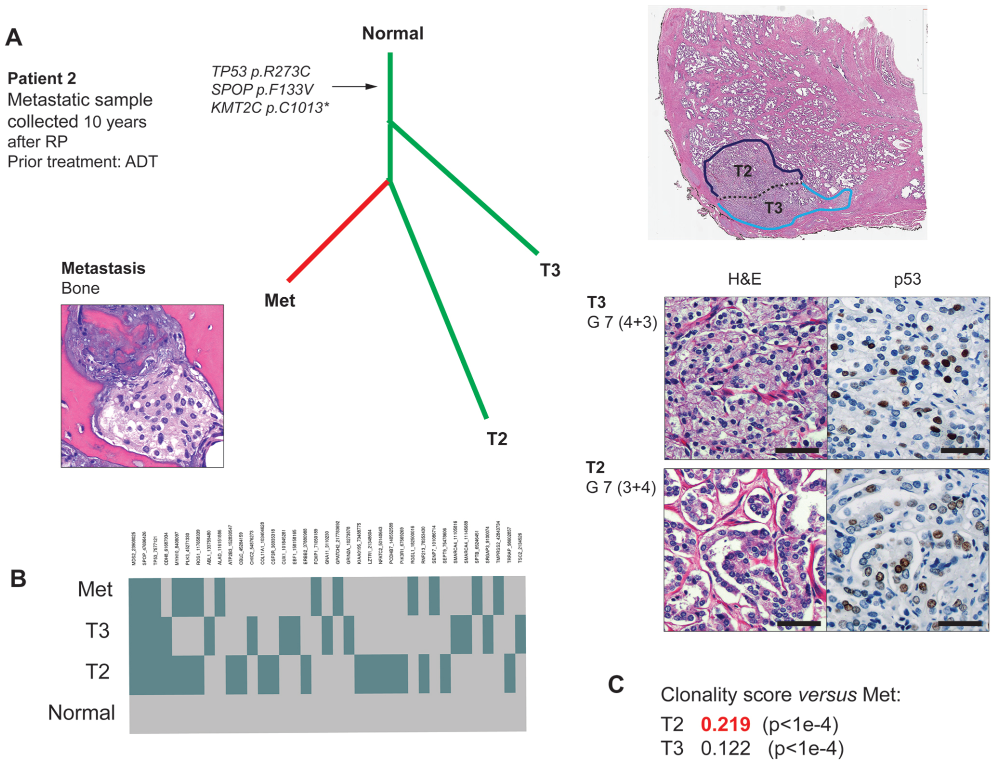


**Supplementary figure S2.** A summary of pathology features, phylogeny and clonality analyses for Patient 2. **A.** Phylogenetic reconstruction using 37 genes and 47 events and pathology features corresponding to each focus. Histomorphology and p53 immunostaining are shown. **B.** A schematic heatmap representation of SNV events used in the phylogenetic reconstruction across samples. **C.** A summary of clonality score results. RP: radical prostatectomy; ADT: androgen deprivation therapy; H&E: hematoxylin-eosin. Scale bars: 50 μm.

### Supplementary figure S3

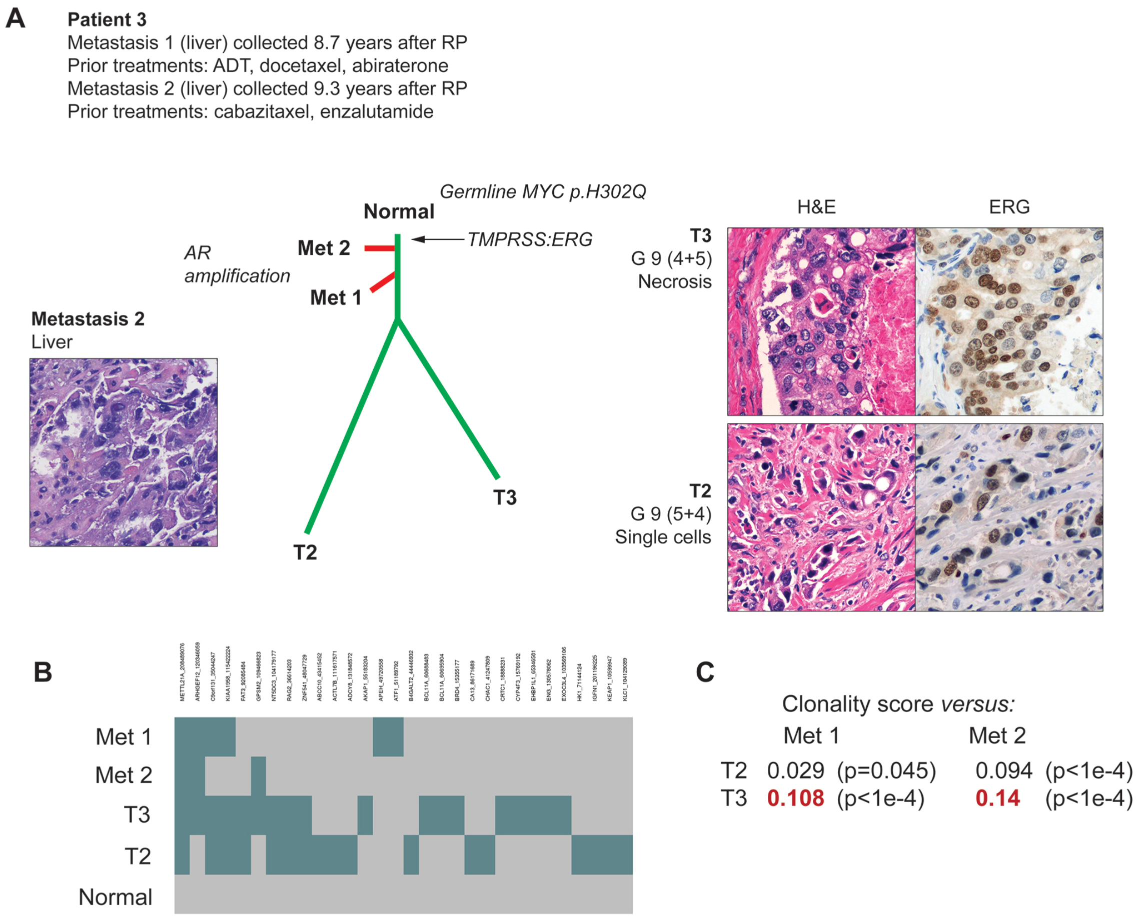


**Supplementary figure S3.** A summary of pathology features, phylogeny and clonality analyses for Patient 3. **A.** Phylogenetic reconstruction using 50 genes and 64 events and pathology features corresponding to each focus. Histomorphology and ERG immunostaining are shown. **B.** A schematic heatmap representation of the most significant SNV events used in the phylogenetic reconstruction across samples. **C.** A summary of clonality score results. RP: radical prostatectomy; ADT: androgen deprivation therapy; H&E: hematoxylin-eosin.

### Supplementary figure S4

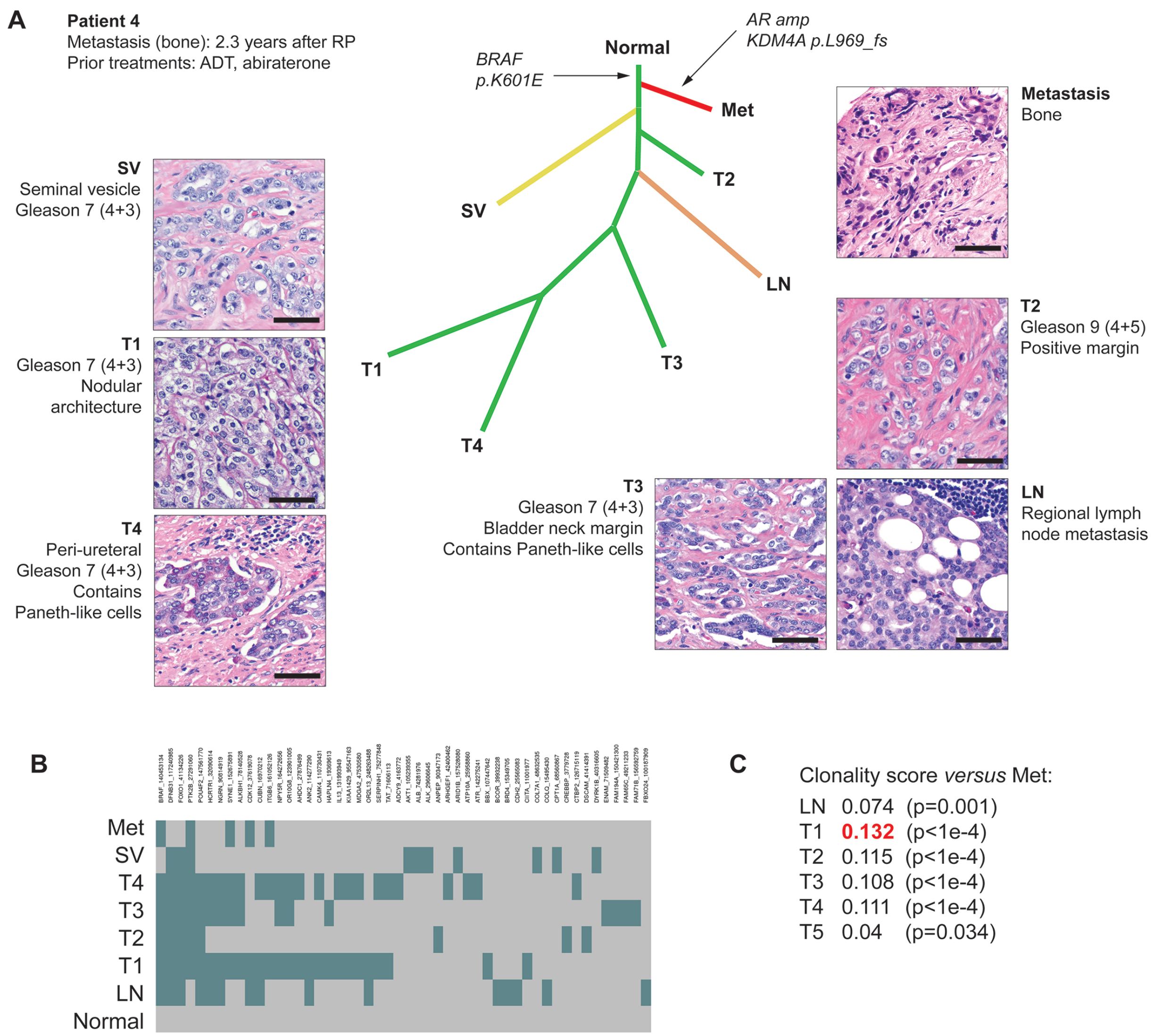


**Supplementary figure S4.** A summary of pathology features, phylogeny and clonality analyses for Patient 4. **A.** Phylogenetic reconstruction using 87 genes and 142 events and pathology features corresponding to each focus. **B.** A schematic heatmap representation of the most significant SNV events used in the phylogenetic reconstruction across samples. **C.** A summary of clonality score results. RP: radical prostatectomy; ADT: androgen deprivation therapy. Scale bars: 50 μm.

### Supplementary figure S5

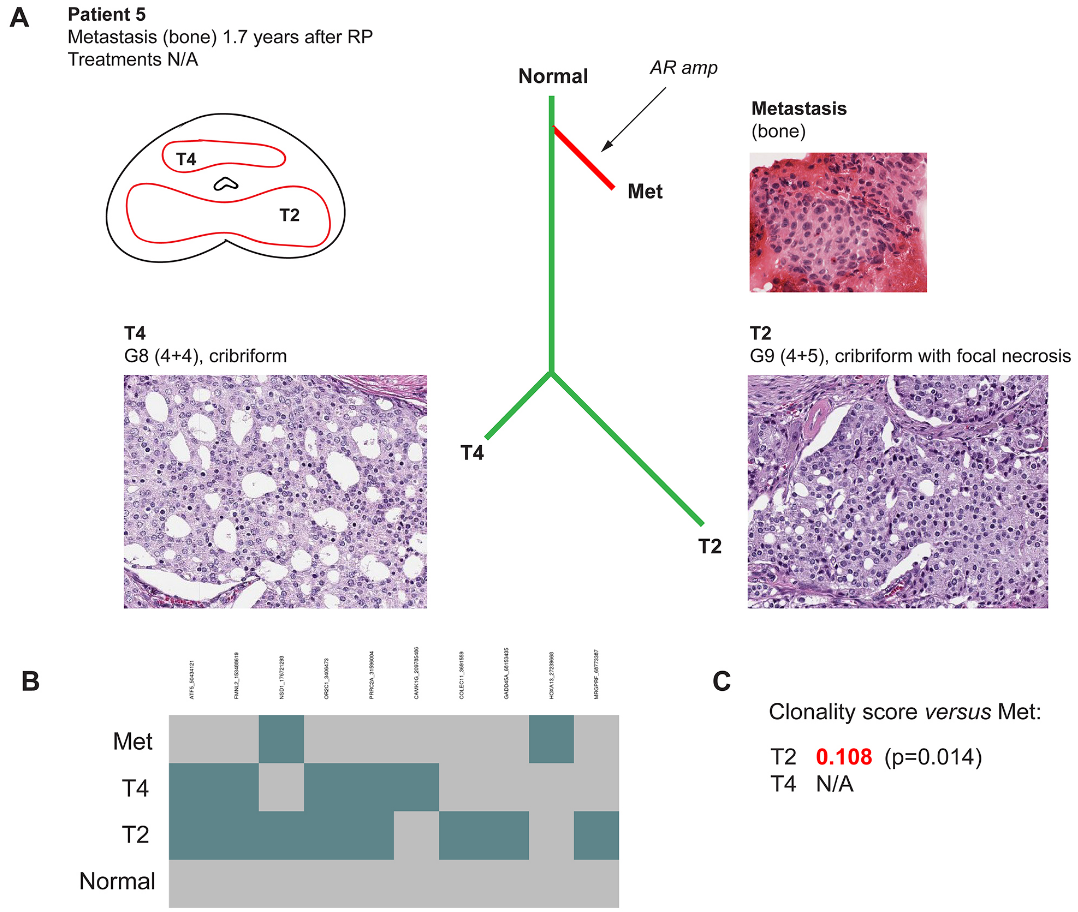


**Supplementary figure S5.** A summary of pathology features, phylogeny and clonality analyses for Patient 5. **A.** Phylogenetic reconstruction using 10 genes and 15 events and pathology features corresponding to each focus. **B.** A schematic heatmap representation of SNV events used in the phylogenetic reconstruction across samples. The clonality scores could not be computed for this case. RP: radical prostatectomy; N/A: not available.

### Supplementary figure S6

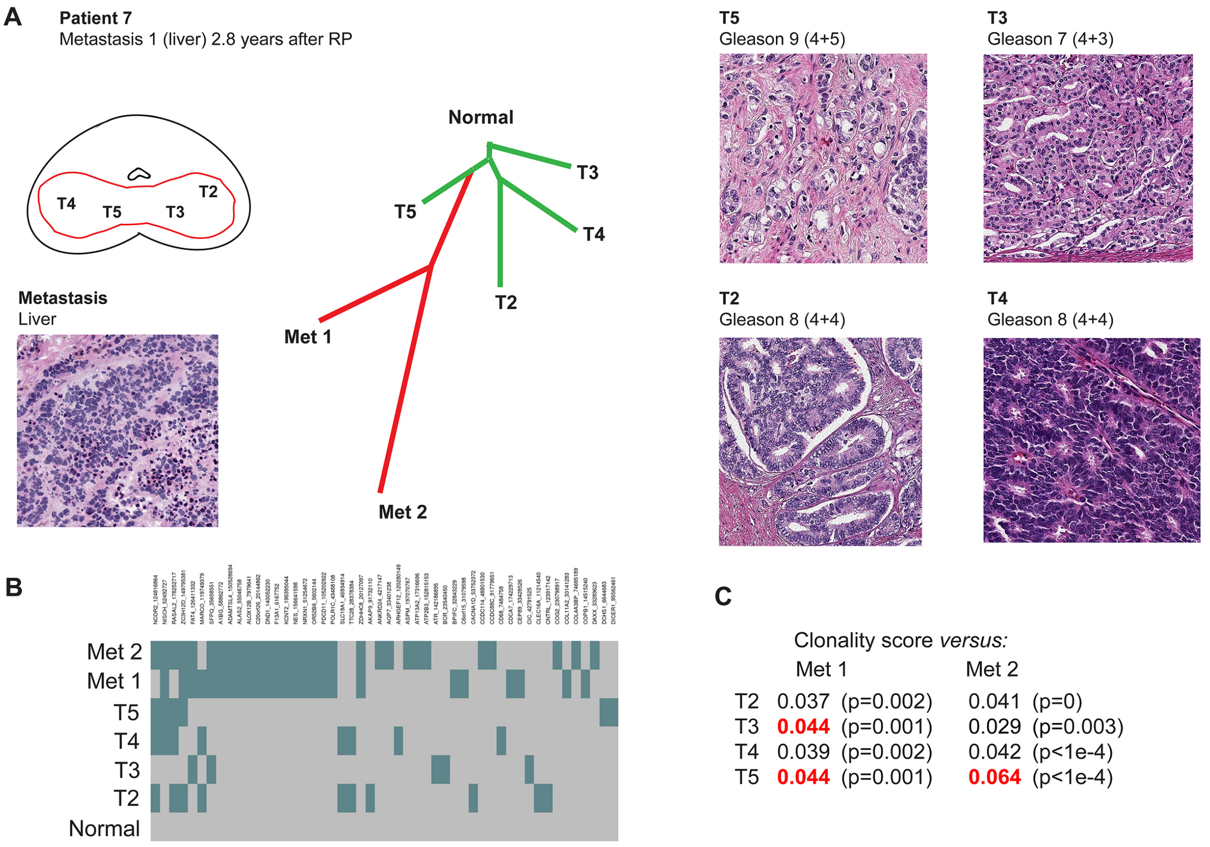


**Supplementary figure S6.** A summary of pathology features, phylogeny and clonality analyses for Patient 7. **A.** Phylogenetic reconstruction using 111 genes and 145 events and pathology features corresponding to each focus. **B.** A schematic heatmap representation of the most significant SNV events used in the phylogenetic reconstruction across samples. **C.** A summary of clonality score results. RP: radical prostatectomy.

**Supplementary figure S7**

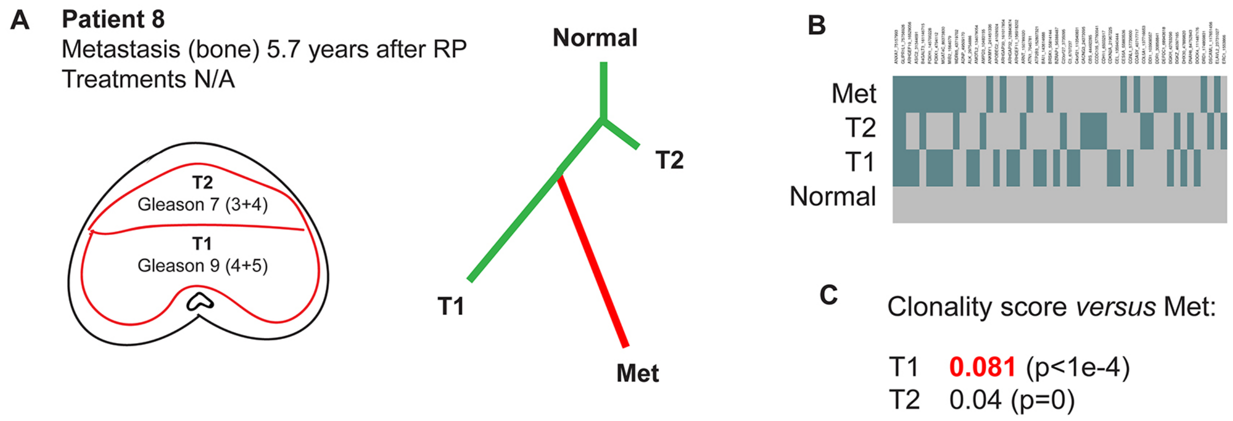


**Supplementary figure S7.** A summary of pathology features, phylogeny and clonality analyses for Patient 8. **A.** Phylogenetic reconstruction using 10 genes and 15 events and a schematic representation of the two tumor foci. **B.** A schematic heatmap representation of SNV events used in the phylogenetic reconstruction across samples. **C.** A summary of clonality score results.

### Supplementary figure S8

**A**

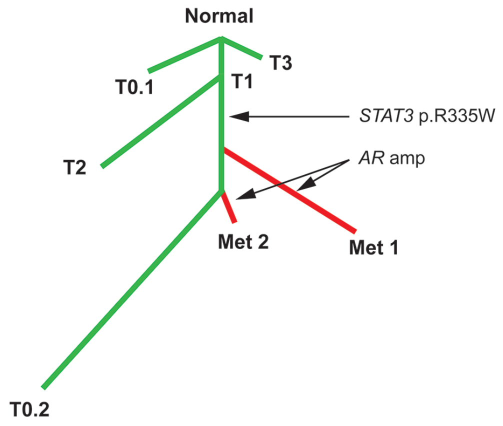


**B**

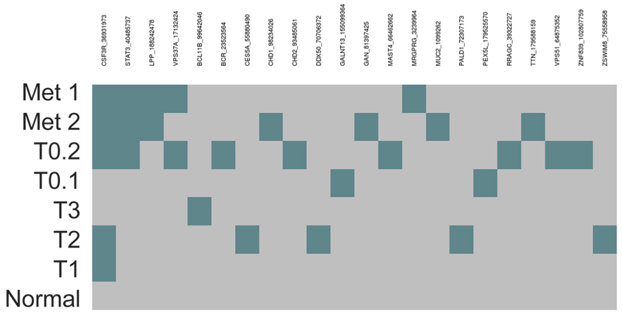


**C**

| | Clonality score *versus:* | | | |
|---|---|---|---|---|
| | Met 1 | | Met 2 | |
| T1 | 0.12 | (p=0.011) | 0.197 | (p=0.007) |
| T2 | 0.088 | (p=0.016) | 0.108 | (p=0.014) |
| T3 | NA | | NA | |
| T0.1 | NA | | NA | |
| T0.2 | **0.14** | (p<1e-4) | **0.27** | (p<1e-4) |

**Supplementary figure S8.** A summary of pathology features, phylogeny and clonality analyses for Patient 10. **A.** Phylogenetic reconstruction using 22 genes and 30 events and pathology features corresponding to each focus. **B.** A schematic heatmap representation of SNV events used in the phylogenetic reconstruction across samples. **C.** A summary of clonality score results.

**Supplementary figure S9**

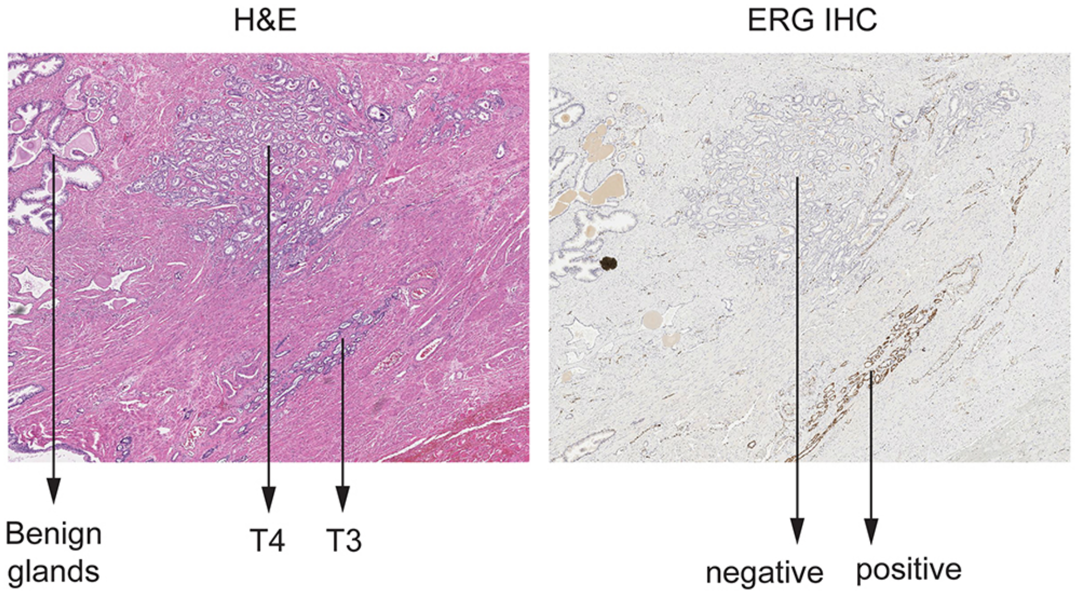


**Supplementary figure S9.** Patient 6, primary tumor focus T4. This focus showed some *TMPRSS2-ERG* signal on RNA-seq, but was negative for ERG by IHC. The topographic proximity of the T3 focus (ERG-positive) and contamination during the macrodissection step may have accounted for this discrepancy. H&E: hematoxylin eosin, IHC: immunohistochemistry.
